# NuSAP1 promotes bipolar spindle assembly in *Trypanosoma brucei* by bundling spindle microtubules

**DOI:** 10.1101/2025.08.25.672125

**Authors:** Qing Zhou, Yasuhiro Kurasawa, Thiago Souza Onofre, Ziyin Li

## Abstract

The parasitic protozoan *Trypanosoma brucei* assembles a bipolar mitotic spindle and undergoes a closed mitosis to segregate its megabase chromosomes and mini-chromosomes through mechanisms that are distinct from its mammalian host. This parasite employs a subset of trypanosome-specific nucleus- and spindle-associated proteins (NuSAPs) to regulate mitosis, but the mechanistic roles of these proteins remain poorly understood. Here, we performed biochemical and molecular characterization of NuSAP1 and analyzed the functional interplay of NuSAP1 with its interacting and proximal proteins. NuSAP1 localizes to the mitotic spindle with spindle pole enrichment, and interacts with the spindle-associated and spindle pole-enriched proteins NuSAP4 and SPB1 through distinct structural motifs. NuSAP1 and NuSAP4 are interdependent for protein stability, and NuSAP1 is required for SPB1 localization. Further, NuSAP1 bundles microtubules *in vitro*, and depletion of NuSAP1 disrupts bipolar spindle assembly. Finally, knockdown of NuSAP1 disrupts the localization of its proximal proteins MAP103 and TbMlp2 to spindle poles. Together, these results uncover the mechanistic role of NuSAP1 in bipolar spindle assembly by bundling spindle microtubules and promoting spindle pole complex formation, underscoring unusual regulatory mechanisms for mitosis in this early divergent unicellular eukaryote.

## Introduction

*Trypanosoma brucei* is an early branching parasitic protozoan causing sleeping sickness in humans and nagana in cattle and poses severe threats to public health and economy in sub-Saharan African. This parasite alternates between the insect vector and the mammalian host by transforming between the insect-infective procyclic form and the mammal-infective bloodstream form, which exhibit distinct morphological and biological features. Within the insect midgut and the mammalian bloodstream, this unicellular eukaryote proliferates and divides through binary fission along its longitudinal axis by an actomyosin-independent mechanism (Hammarton et al., 2007). Despite the unusual mechanism of cytokinesis, however, *T. brucei* employs a typical eukaryotic cell cycle control system, consisting the DNA replication phase (S phase), the mitotic phase (M phase), and two gap phases (G1 and G2 phases) that prepare the cell for chromosome replication and segregation, respectively, and a conserved cyclin/cyclin-dependent kinase machinery that governs the progression of the cell cycle (Li, 2012). Distinctive features in cell cycle control between the procyclic and the bloodstream forms(Hammarton, 2007) and between *T. brucei* and its human host (Li, 2012) have been observed, highlighting the potential exploitation of cell cycle regulatory proteins as drug targets.

*T. brucei* undergoes a closed mitosis without breaking down its nuclear envelope during mitosis and assembles an intra-nuclear spindle with fewer microtubule fibers than the spindle in its mammalian host (Ogbadoyi et al., 2000). Spindle assembly in *T. brucei* does not appear to require the γ-tubulin ring complex, the evolutionarily conserved microtubule nucleation machinery (McKean et al., 2003; Zhou and Li, 2015), and there is no evidence to support the existence of a canonical centriole structure at the spindle pole (Ogbadoyi et al., 2000), suggesting a likely chromatin-based spindle assembly mechanism in trypanosomes. *T. brucei* lacks the homolog of CENP-A, the centromere-specific histone H3 variant and a hallmark of the kinetochore which specifies the kinetochore assembly site on chromosomes (Hori and Fukagawa, 2012), and assembles kinetochores with highly divergent proteins (Akiyoshi and Gull, 2014; D’Archivio and Wickstead, 2017; Nerusheva and Akiyoshi, 2016). *T. brucei* also lacks homologs of numerous evolutionarily conserved mitotic regulators, including the spindle motor protein BimC homolog, the kinetochore motor protein CENP-E homolog, and the spindle-assembly checkpoint protein homologs (Berriman et al., 2005). These distinctive features suggest unusual control mechanisms and regulatory pathways governing bipolar spindle assembly and chromosome segregation in *T. brucei*.

The spindle apparatus is a microtubule-based macromolecular machine that plays crucial roles in capturing, aligning, and separating the sister chromatids during mitosis through the orchestrated physical forces exerted by spindle-associated motor and non-motor proteins (Petry, 2016). These spindle-associated proteins (SAPs) have diverse molecular structures and biochemical functions, and play roles by promoting bipolar spindle assembly, spindle orientation and positioning, spindle elongation, and microtubule-kinetochore interactions (Petry, 2016). SAPs discovered in *T. brucei* also include motor and non-motor proteins, many of which are kinetoplastid-specific proteins with uncharacterized biochemical functions (Billington et al., 2023). At least four kinesin proteins have been identified as SAPs in *T. brucei*, including two orphan kinesins, KIN-A and KIN-B (Li et al., 2008), which target the Aurora B kinase homolog TbAUK1 (Ballmer and Akiyoshi, 2024), a kinesin-13-family kinesin named Kif13-1/KIN13-1 that depolymerizes spindle microtubules and regulates spindle dynamics (Chan et al., 2010; Wickstead et al., 2010), and an orphan kinesin named KIN-F whose function remains to be explored (Zhou et al., 2018). Other SAPs include the nuclear pore complex protein TbMlp2/TbNup92, which associates with spindle poles and promotes chromosome segregation (Holden et al., 2014; Morelle et al., 2015), two microtubule-severing enzymes fidgetin (Zhou et al., 2018) and spastin, the latter of which regulates spindle dynamics and chromosome segregation (Souza Onofre et al., 2025), four kinetoplastid-specific, nucleus- and spindle-associated proteins (NuSAPs), which are required for chromosome segregation (Zhou et al., 2018; Zhou and Li, 2024), and the spindle-localized and spindle pole-enriched proteins SPB1 (Zhou et al., 2018) and MAP103 (Hayashi and Akiyoshi, 2018), whose functions remain elusive. With the exception of NuSAP3, which maintains the stability of its interacting partner protein Kif13-1/KIN13-1 (Zhou et al., 2018), however, the mechanistic roles of the other three NuSAPs have not yet been elucidated.

In this report, we characterized the role of NuSAP1 in spindle assembly and chromosome segregation in the procyclic (insect) form of *T. brucei* by biochemical and molecular and cell biological approaches. We demonstrated that NuSAP1 bundles microtubules *in vitro* and promotes bipolar spindle assembly in trypanosome cells. We also showed that NuSAP1 interacts with NuSAP4 and SPB1 through distinct structural domains and regulates the stability of NuSAP1 and the localization of SPB1. This work uncovered the mechanistic role of NuSAP1 in regulating spindle assembly and underscored the unusual control mechanisms for spindle assembly by kinetoplastid-specific SAPs in *T. brucei*.

## Results

### NuSAP1 is a kinetoplastid-specific SAP that bundles microtubules *in vitro*

Using AlphaFold3, we predicted that NuSAP1 contains an N-terminal intrinsically disordered region (IDR1), a long coiled-coil motif (CC1) of ∼640 a.a., another intrinsically disordered region (IDR2), and a short coiled-coil motif (CC2) of ∼150 a.a. (Fig. 1A). Co-immunofluorescence microscopy with endogenous PTP-tagged NuSAP1 and 3HA-tagged β-tubulin showed that NuSAP1 localizes to the entire spindle at metaphase and is additionally enriched at pindle poles thereafter until anaphase B, during which NuSAP1 also starts to spread into the nucleus (Fig. 1B). The association of NuSAP1 with the spindle during metaphase and the enrichment of NuSAP1 at spindle poles during anaphase were further confirmed by co-immunostaining with the spindle-associated protein Kif13-1/KIN13-1 and the spindle pole-localized protein TbMlp2 (Fig. 1C).

**Figure 1.**
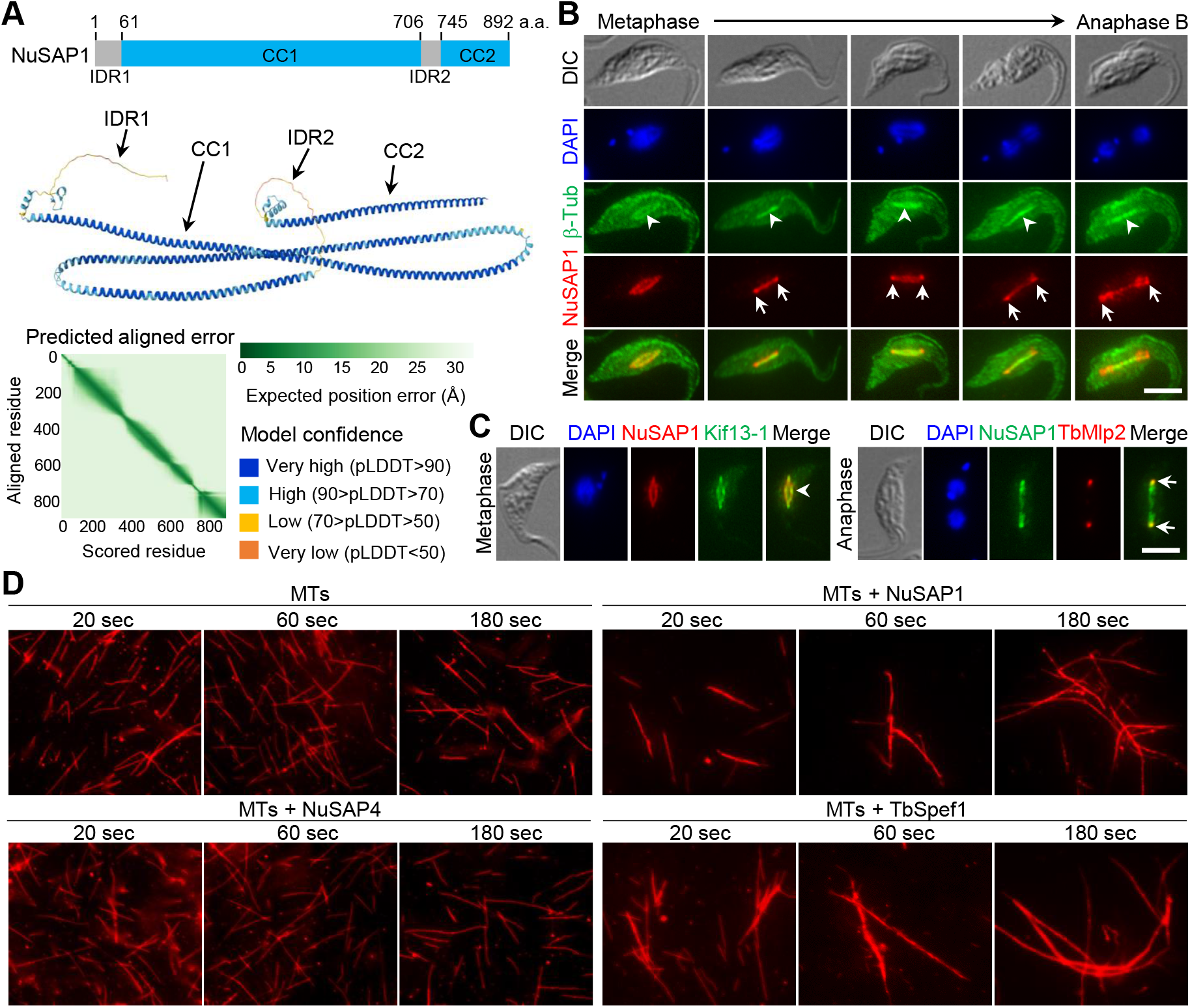
NuSAP1 is a spindle-associated protein that bundles microtubules *in vitro*. (**A**). Schematic illustration of NuSAP1 and its structural domains. IDR: intrinsically disordered region; CC: coiled coil. (**B**). NuSAP1 structure predicted by AlphaFold3. (**C**). Subcellular localization of NuSAP1 from metaphase to late anaphase in procyclic trypanosomes. Endogenous NuSAP1-PTP and 3HA-β-tubulin were detected by anti-Protein A pAb and anti-HA mAb, respectively. Open arrowheads indicate the spindle, and arrows indicate NuSAP1 enrichment at spindle poles. Scale bar: 5 μm. (**D**). Co-localization of NuSAP1 with the spindle marker protein Kif13-1 and the spindle pole marker protein TbMlp2. Endogenous NuSAP1-PTP and TbMlp2-3HA were immunostained by anti-Protein A pAb and Kif13-1-3HA was immunostained by anti-HA mAb. Open arrowhead indicates the spindle, and arrows indicate spindle poles. Scale bar: 5 μm. (**E**). *In vitro* microtubule-bundling assay using purified recombinant NuSAP1, NuSAP4, and TbSpef1 proteins. Microtubules (MTs) were assembled *in vitro* and then incubated with purified recombinant proteins for up to 3 minutes.

The association of NuSAP1 with the spindle prompted us to test the potential role of NuSAP1 in regulating microtubule dynamics. By incubating purified recombinant NuSAP1 with *in vitro* assembled microtubules, we found that NuSAP1 promoted microtubule bundle formation (Fig. 1D). As a positive control, purified recombinant TbSpef1 also promoted microtubule bundling under same experimental conditions (Fig. 1D). However, purified recombinant NuSAP4 exerted no detectable effects on microtubules (Fig. 1D). Because NuSAP1 localizes to the spindle, this result suggests that NuSAP1 may promote spindle microtubule bundling in trypanosome cells.

### NuSAP1 is required for bipolar spindle assembly

We previously reported that the mitotic spindle was still detectable in NuSAP1 RNAi cells (Zhou et al., 2018), but whether the spindle structure was affected by NuSAP1 RNAi was not investigated. To test whether NuSAP1 RNAi may disrupt spindle structure, we performed immunofluorescence microscopy with the anti-β-tubulin antibody KMX-1 and two spindle-associated proteins, MAP103 and KIN-F, as spindle markers (Fig. 2). In control cells, the metaphase spindle was detected as either a diamond-shaped structure or a straight line, which was strictly confined within the nucleus (Fig. 2A-C). In NuSAP1 RNAi-induced cells, however, the metaphase spindle either lost the typical diamond shape or was detected as an elongated line extending outside of the DAPI-stained nucleus (Fig. 2A-C). Such malformed metaphase spindles were detected in ∼56% of the metaphase cells after NuSAP1 RNAi for 24 h (Fig. 2D). In control cells, the anaphase spindle was always detected as a straight line confined within the segregating nuclei (Fig. 2A-C). The anaphase spindle appeared to be composed of bundled microtubules or clustered microtubules, as previously shown by electron microscopy (Ogbadoyi et al., 2000). In NuSAP1 RNAi cells, however, the anaphase spindle was either distorted/bent or irregularly shaped, with one pole of the spindle occasionally extended out of the DAPI-stained nucleus (Fig. 2A-C). These malformed spindles were detected in ∼77% of the anaphase cells after NuSAP1 RNAi induction for 24 h (Fig. 2D). Together, these observations suggest that NuSPA1 is required for proper assembly of a bipolar spindle in *T. brucei*, likely by promoting spindle microtubule bundling, as suggested by its *in vitro* microtubule-bundling activity (Fig. 1D).

**Figure 2.**
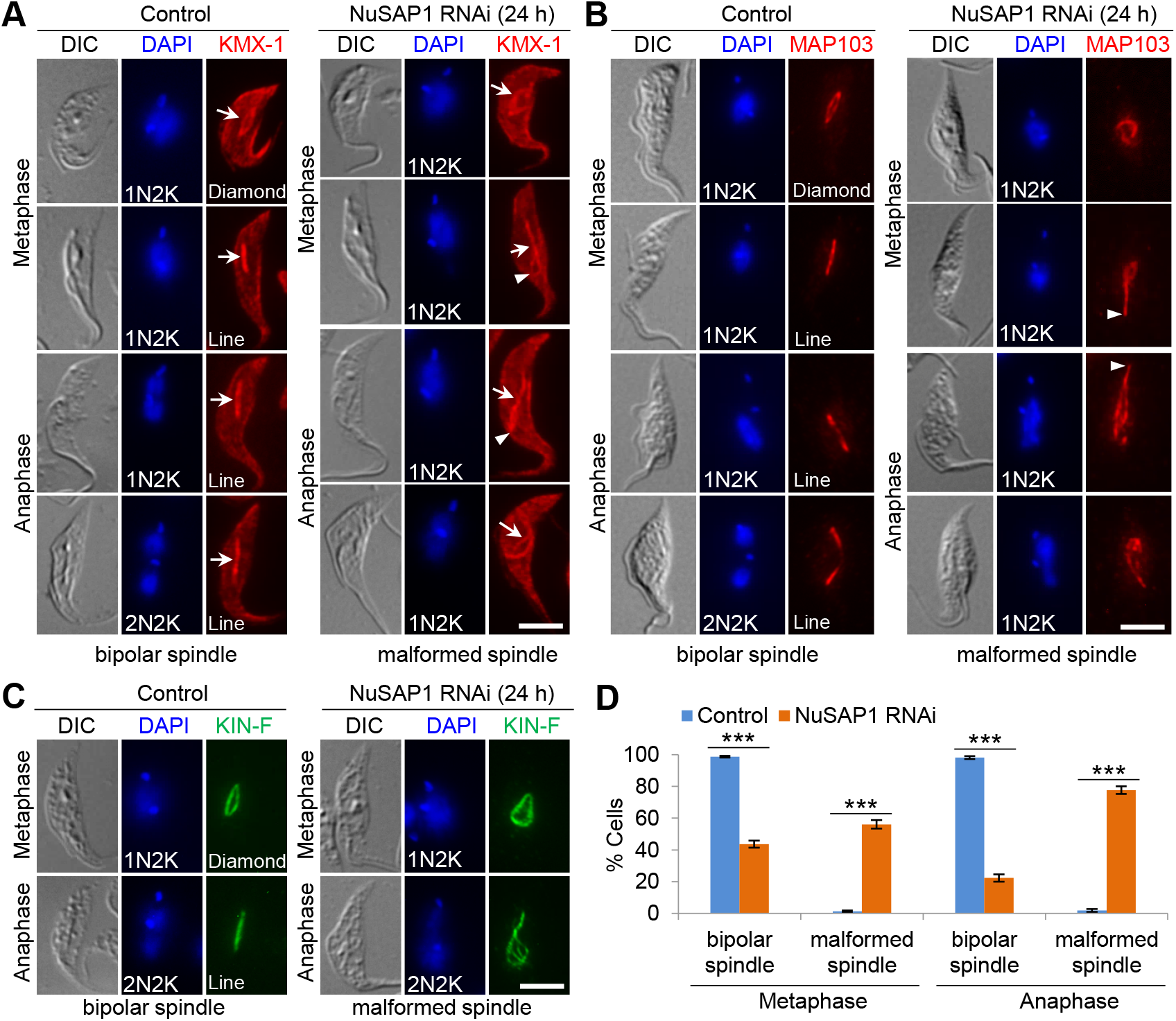
NuSAP1 is required for bipolar spindle assembly. (**A**). Effect of NuSAP1 RNAi on spindle assembly using β-tubulin as a spindle marker. KMX-1 antibody was used to detect β-tubulin. Arrows indicate spindle and arrowheads indicate the spindle pole that was extended outside of the DAPI-stained nucleus. Scale bar: 5 μm. (**B**). Effect of NuSAP1 RNAi on spindle assembly using the spindle-associated protein MAP103 as a spindle marker. Endogenous MAP103-PTP was detected by anti-Protein A pAb. Arrowheads indicate the spindle pole extended outside of the nuclear DNA. Scale bar: 5 μm. (**C**). Effect of NuSAP1 RNAi on spindle assembly using the spindle-associated kinesin protein KIN-F as a spindle marker. Endogenous KIN-F-3HA was detected by anti-HA mAb. Scale bar: 5 μm. (**D**). Quantitation of metaphase and anaphase cells with a malformed spindle in control and NuSAP1 RNAi cells. Error bars indicated S.D. from three independent experiments. ***, *p*<0.001.

### NuSAP1 interacts with NuSAP4 and is required for NuSAP4 stability

We previously performed proximity-dependent protein biotinylation (BioID) using NuSAP1 as bait (Zhou et al., 2018), and identified four spindle-associated or spindle pole-enriched proteins, NuSAP4, SPB1, TbMlp2, and MAP103 (Fig. 3A). Like NuSAP1, NuSAP4 also localizes to the nucleus prior to mitotic onset and then to the spindle during mitosis (Zhou et al., 2018). Thus, we first tested whether NuSAP1 may co-localize with and interact with NuSAP4. Co-immunofluorescence microscopy confirmed their co-localization on the spindle during metaphase and anaphase (Fig. 3B), and co-immunoprecipitation showed that NuSAP4 was able to pull down NuSAP1 from trypanosome cell lysate (Fig. 3C), demonstrating that NuSAP1 and NuSAP4 form a complex *in vivo* in trypanosome cells. Using AlphaFold3, we predicted the structure of the NuSAP1-NuSAP4 complex, which showed that NuSAP4 associates with the C-terminal CC2 motif of NuSAP1 (Fig. 3D, left). A closer examination of the CC2 motif showed that it has an almost identical fold as NuSAP4, despite little sequence similarity, and the predicted CC2^NuSAP1^-NuSAP4 structure (Fig. 3D, bottom right) resembles the predicted NuSAP4 homodimer structure (Zhou and Li, 2024). It is possible that NuSAP4 does not form a homodimer, but instead it forms a heterodimer with NuSAP1 through binding to the CC2 motif of NuSAP1.

**Figure 3.**
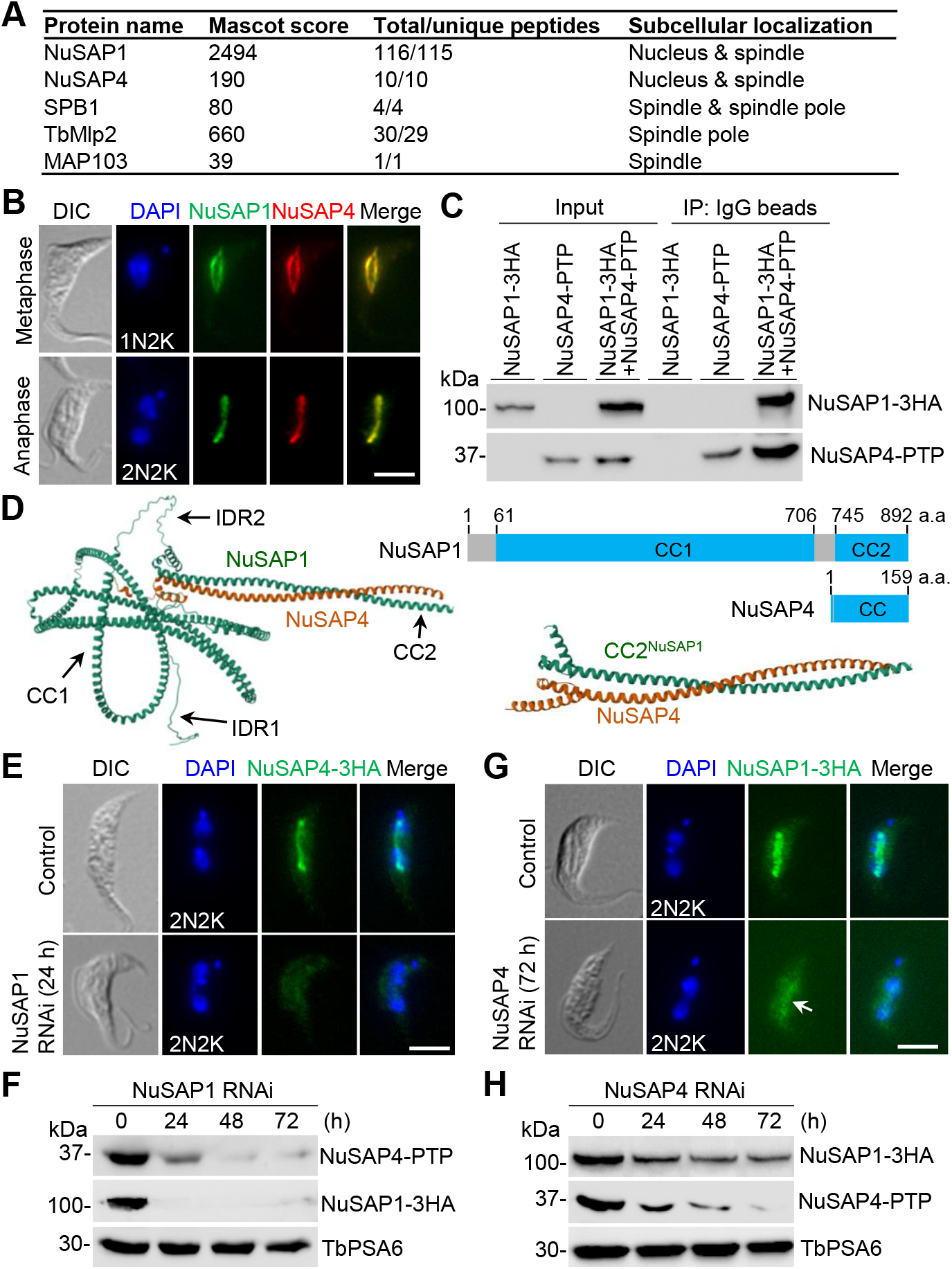
NuSAP1 and NuSAP4 form a complex and are interdependency for protein stability. (**A**). List of NuSAP1-proximal proteins identified by NuSAP1 BioID. (**B**). Co-localization of NuSAP1 and NuSAP4 at the spindle. Endogenous NuSAP1-3HA and NuSAP4-PTP were detected by anti-HA mAb and anti-Protein A pAb, respectively. Scale bar: 5 μm. (**C**). NuSAP1 interacts with NuSAP4 *in vivo* in trypanosomes. NuSAP4-PTP and NuSAP1-3HA were co-immunoprecipitated, and then detected by anti-Protein A pAb and anti-HA mAb, respectively. IP: immunoprecipitation. (**D**). NuSAP1-NuSAP4 hetero-dimer structure and CC2^NuSAP1^-NuSAP4 hetero-dimer structure predicted by AlphaFold3. The top right panel shows the schematic drawing of NuSAP1 and NuSAP4 structural domains. (**E**). Effect of NuSAP1 RNAi on the localization of NuSAP4. NuSAP4-3HA was detected by anti-HA mAb. Scale bar: 5 μm. (**F**). Effect of NuSAP1 RNAi on NuSAP4 stability. Endogenous NuSAP4-PTP and NuSAP1-3HA were detected by anti-Protein A pAb and anti-HA mAb, respectively. TbPSA6 served as a loading control. (**G**). Effect of NuSAP4 RNAi on the localization of NuSAP1. NuSAP1-3HA was detected by anti-HA mAb. Scale bar: 5 μm. (**H**). Effect of NuSAP4 RNAi on NuSAP1 stability. Endogenous NuSAP1-3HA and NuSAP4-PTP were detected by anti-HA mAb and anti-Protein A pAb, respectively. TbPSA6 served as a loading control.

We next investigated the functional interplay between NuSAP1 and NuSAP4 in trypanosome cells. Knockdown of NuSAP1 caused the disappearance of NuSAP4 signal from the spindle, shown by immunofluorescence microscopy (Fig. 3E) and a drastic reduction of NuSAP4 protein level, shown by western blotting (Fig. 3F). Reciprocally, knockdown of NuSAP4 reduced the NuSAP1 signal intensity from the spindle (Fig. 3G) and moderately reduced the NuSAP1 protein level (Fig. 3H). Thus, NuSAP1 and NuSAP4 are interdependent for their stability, albeit that RNAi of NuSAP4 exerted weaker effects than RNAi of NuSAP1.

### The CC2 motif of NuSAP1 is required for the function of NuSAP1 and the interaction with NuSAP4

To test the potential requirement of the CC2 motif for the cellular function of NuSAP1 and the interaction with NuSAP4, we performed RNAi complementation experiments with the CC2-deletion mutant of NuSAP1 and, as a control, the wild-type NuSAP1. We first generated a NuSAP1-3’UTR RNAi cell line by targeting against the 3’UTR of the *NuSAP1* gene, and western blotting confirmed the moderate knockdown efficiency (Fig. 4A). This NuSAP1-3’UTR RNAi cell line showed strong growth defects (Fig. 4B), similar to the NuSAP1 RNAi cell line that targets the *NuSAP1* coding region (Zhou et al., 2018). Ectopic overexpression (OE) of wild-type NuSAP1 in NuSAP1-3’UTR RNAi cells (hereafter referred to as NuSAP1-3’UTR RNAi/NuSAP1 OE) partially restored the growth defects, but ectopic overexpression of NuSAP1-ΔCC2 in NuSAP1-3’UTR RNAi cells failed to rescue the growth defects (Fig. 4B), suggesting the loss of function of the NuSAP1-ΔCC2 mutant.

**Figure 4.**
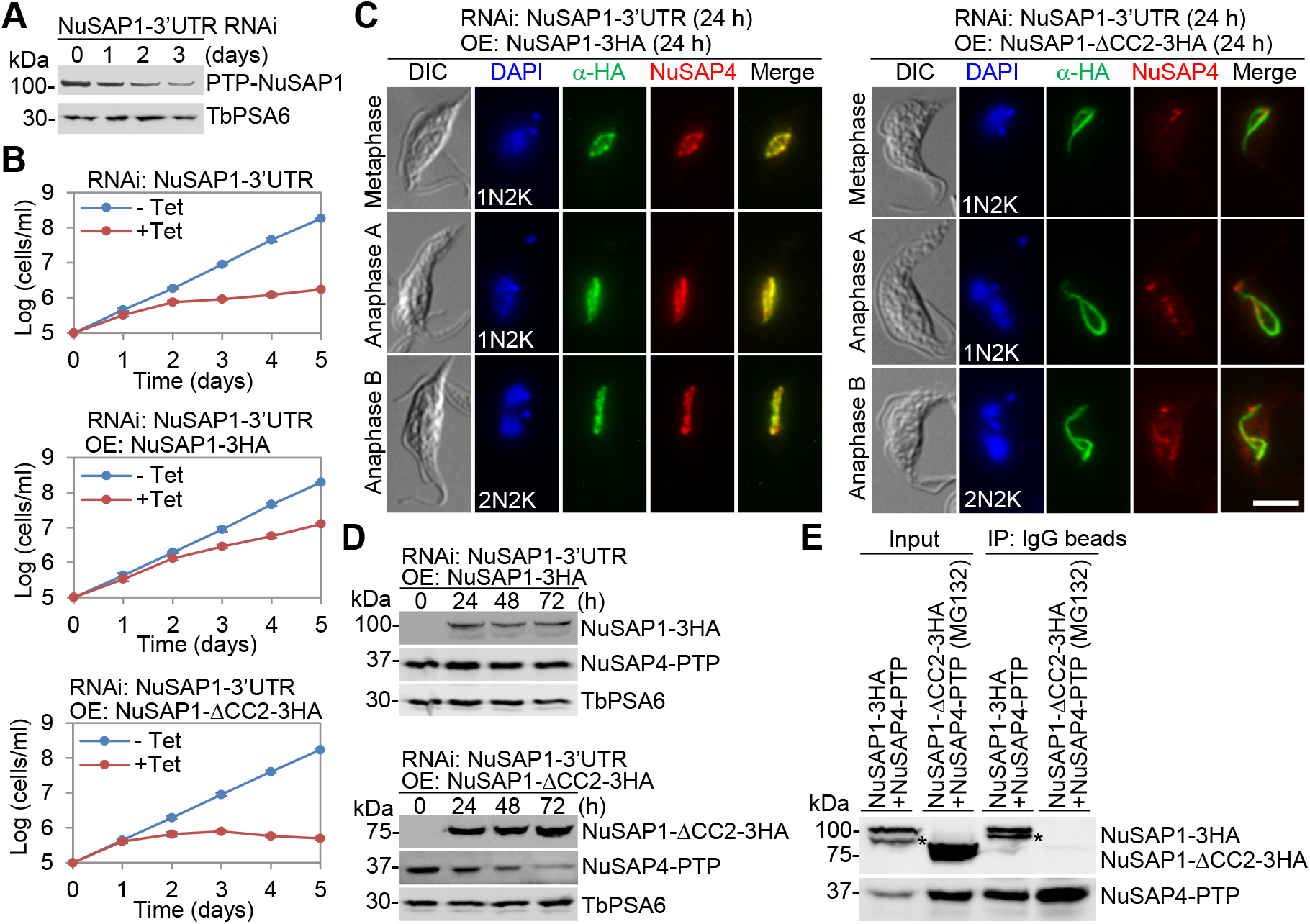
The CC2 motif is required for the function of NuSAP1 and the interaction with NuSAP4. (**A**). Western blotting to detect NuSAP1 protein level after RNAi. NuSAP1 was endogenously tagged with an N-terminal PTP and detected by anti-Protein A antibody. TbPSA6 served as a loading control. (**B**). Growth curves of NuSAP1-3’UTR RNAi cell line and its complementation cell lines expressing NuSAP1 or NuSAP1-ΔCC2. (**C**). Effect of the expression of NuSAP1-ΔCC2 on NuSAP4 localization. Expression of the full-length NuSAP1 served as a control. Scale bar: 5 μm. (**D**). Effect of the expression of NuSAP1-ΔCC2 on NuSAP4 stability. Expression of the full-length NuSAP1 served as a control. (**E**). Co-immunoprecipitation to test the interaction between NuSAP1-ΔCC2 and NuSAP4. The full-length NuSAP1 served as a control.

We next endogenously tagged NuSAP4 with a PTP epitope in these two RNAi complementation cell lines and examined the localization of NuSAP1, NuSAP1-ΔCC2, and NuSAP4 by co-immunofluorescence microscopy (Fig. 4C). The ectopically overexpressed NuSAP1 and NuSAP1-ΔCC2 both localized to the spindle, despite the abnormal spindle shape in the NuSAP1-3’UTR RNAi/NuSAP1-ΔCC2 OE cells (Fig. 4C). In NuSAP1-3’UTR RNAi/NuSAP1 OE cells, NuSAP4 co-localized with the ectopically overexpressed NuSAP1 (Fig. 4C); however, in NuSAP1-3’UTR RNAi/NuSAP1-ΔCC2 OE cells, NuSAP4 signal was weaker and was detected as punctate dots along the abnormal-shaped spindle (Fig. 4C), suggesting the disruption of NuSAP4 localization by deleting the CC2 motif in NuSAP1. Western blotting showed that the NuSAP4 protein level was unaffected in NuSAP1-3’UTR RNAi/NuSAP1 OE cells, but it was gradually reduced in NuSAP1-3’UTR RNAi/NuSAP1-ΔCC2 OE cells (Fig. 4D), consistent with the lower NuSAP4 fluorescence intensity in these cells (Fig. 4C). Co-immunoprecipitation showed that the endogenously expressed NuSAP4 was able to pull down ectopically overexpressed NuSAP1, but not NuSAP1-ΔCC2 (Fig. 4E), demonstrating that the CC2 motif is required for NuSAP1 interaction with NuSAP4, in agreement with the AlphaFold3-predicted binding of NuSAP4 to the CC2 domain of NuSAP1. The result also suggests that the interaction with NuSAP1 maintains NuSAP4 stability.

### NuSAP1 interacts with SPB1 and is required for SPB1 localization

The enrichment of NuSAP1 at spindle poles (Fig. 1B) and the identification of SPB1 as a NuSAP1-proximal protein (Fig. 3A) prompted us to test whether NuSAP1 interacts with SPB1. Co-immunofluorescence microscopy showed that NuSAP1 and SPB1 co-localized at spindle poles from metaphase to anaphase B (Fig. 5A), and co-immunoprecipitation showed that NuSAP1 was able to pull down SPB1 from trypanosome cell lysate (Fig. 5B), demonstrating that NuSAP1 and SPB1 form a complex *in vivo*. Since NuSAP4 also interacts with SPB1(Zhou and Li, 2024), NuSAP1, NuSAP4, and SPB1 may form a tri-protein complex. Using AlphaFold3, we predicted the structure of the NuSAP1-NuSAP4-SPB1 complex, which showed that NuSAP4 associates with the CC2 motif of NuSAP1 and SPB1 binds to the IDR2 of NuSAP1 (Fig. 5C).

**Figure 5.**
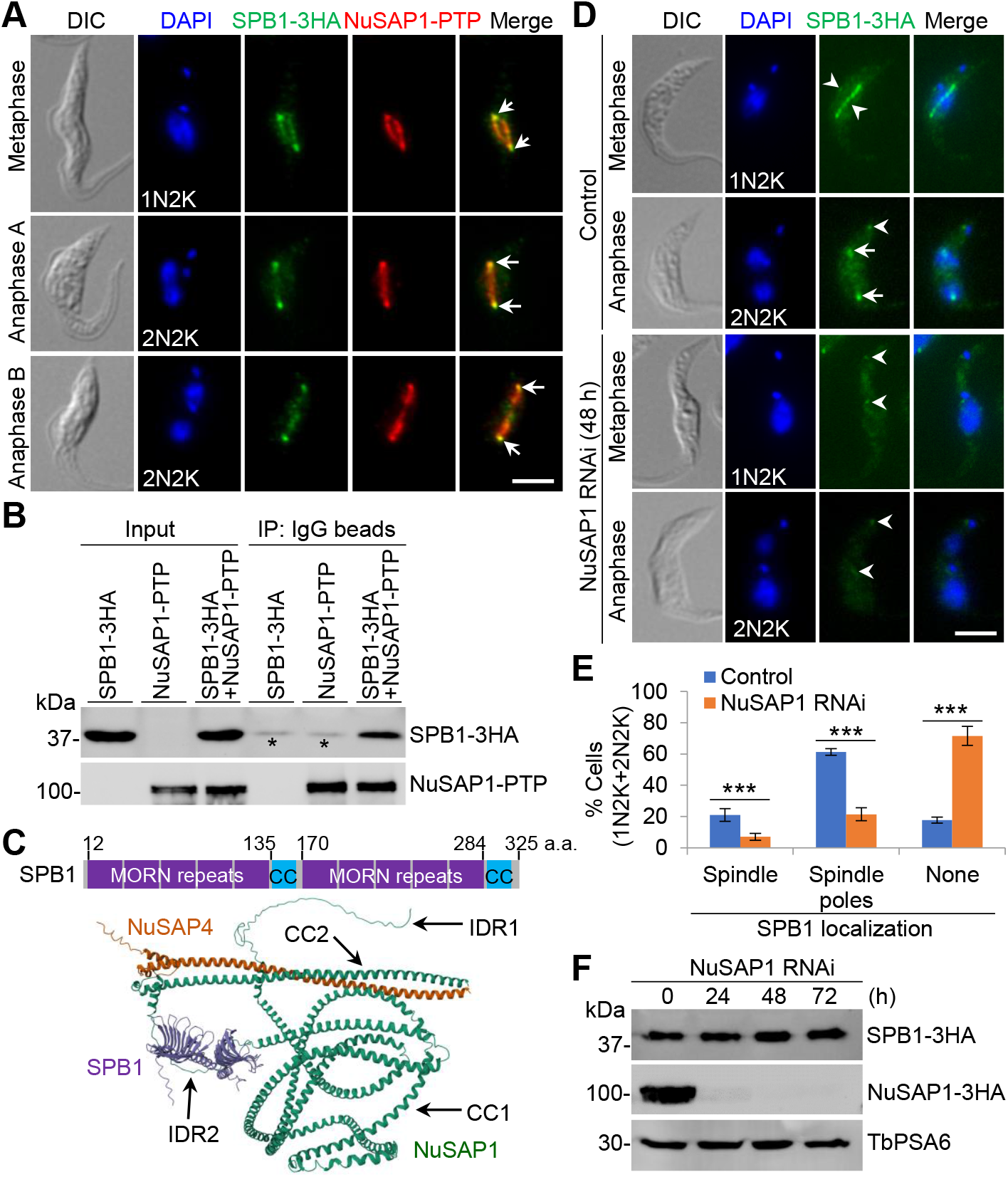
NuSAP1 interacts with SPB1 and is required for SPB1 localization to spindle poles. (**A**). NuSAP1 enriches and co-localizes with SPB1 at the spindle poles. Endogenous NuSAP1-PTP and SPB1-3HA were detected by anti-Protein A pAb and anti-HA mAb, respectively. Arrows indicate the co-localization of NuSAP1 and SPB1 at spindle poles. Scale bar: 5 μm. (**B**). NuSAP1 interacts with SPB1 *in vivo* in trypanosomes. NuSAP1-PTP and SPB1-3HA were co-immunoprecipitated, and then detected by anti-Protein A pAb and anti-HA mAb, respectively. The asterisks indicate a non-specific band. IP: immunoprecipitation. (**C**). Schematic illustration of SPB1 structural domain organization and NuSAP1-NuSAP4-SPB1 hetero-trimer structure predicted by AlphaFold3. (**D**). Effect of NuSAP1 knockdown on SPB1 localization. SPB1 was endogenously tagged with a triple HA epitope and detected by anti-HA mAb. Arrows indicate SPB1-3HA at the spindle poles, whereas arrowheads indicate SPB1-3HA at the basal body. Scale bar: 5 μm. (**E**). Quantitation of cells (1N2K and 2N2K) with different SPB1 localization patterns in control and NuSAP1 RNAi cells. Error bars indicate S.D. from three independent experiments. ***, *p*<0.001. (**F**). Western blotting to detect SPB1 protein levels in control and NuSAP1 RNAi cells. Endogenous SPB1 and NuSAP1, each of which tagged with a triple HA epitope, were detected by anti-HA mAb. TbPSA6 served as a loading control.

We asked whether NuSAP1 knockdown affects SPB1 localization and/or stability, as it does to NuSAP4 (Fig. 4). In control cells, SPB1 was either localized to the entire spindle or was enriched at spindle poles during metaphase and was enriched at spindle poles during anaphase (Fig. 5A, D). Weak SPB1 signal at basal bodies was occasionally detected in some of the cells (Fig. 5D), as reported previously (Billington et al., 2023). In NuSAP1 RNAi cells induced for 48 h, SPB1 localization to the spindle and spindle poles, but not basal bodies, was impaired (Fig. 5D). The 1N2K and 2N2K cells with detectable SPB1 signal at the spindle or spindle poles were reduced from ∼21% to ∼7% and from ∼61% to ∼21%, respectively, and correspondingly, those cells without detectable SPB1 signal were increased from ∼18% to ∼72% (Fig. 5E). Western blotting showed that SPB1 protein levels were not affected in NuSAP1 RNAi cells (Fig. 5F). Thus, NuSAP1 is required for SPB1 localization, but not SPB1 stability. This effect is similar to that of NuSAP4 RNAi on SPB1 (Zhou and Li, 2024).

### The IDR2 domain of NuSAP1 is required for NuSAP1 function

Since AlphaFold3 predicted that SPB1 interacts with the IDR2 of NuSAP1 (Fig. 5C), we tested whether IDR2 is required for the cellular function of NuSAP1 and the interaction with SPB1. To this end, we ectopically overexpressed NuSAP1-ΔIDR2 in the NuSAP1-3’UTR RNAi cell line and found that NuSAP1-ΔIDR2 was unable to rescue the growth defects of the NuSAP1-3’UTR RNAi cells (Fig. 6A), suggesting that IDR2 is essential for NuSAP1 function. To test whether deletion of IDR2 disrupts NuSAP1 interaction with SPB1, we tagged SPB1 with a PTP epitope at its endogenous locus in the NuSAP1-3’UTR RNAi/NuSAP1-ΔIDR2 OE cells and the NuSAP1-3’UTR RNAi/NuSAP1 OE cells. Co-immunoprecipitation showed that endogenously expressed SPB1 was able to pull down ectopically overexpressed NuSAP1, but not NuSAP1-ΔIDR2 (Fig. 6B), demonstrating that IDR2 is required for SPB1 interaction with NuSAP1, consistent with the AlphaFold3 prediction that SPB1 binds to the IDR2 of NuSAP1 (Fig. 5C). We next examined whether the disruption of NuSAP1-SPB1 interaction by deleting IDR2 may affect SPB1 localization by immunofluorescence microscopy. In NuSAP1-3’UTR RNAi/NuSAP1-ΔIDR2 OE cells, SPB1 localization to the spindle and spindle poles was reduced from ∼19% to ∼4% and from ∼64% to ∼10%, respectively (Figs. 6C, D), despite that the protein level of SPB1 was not affected (Fig. 6E). These results demonstrated that the interaction with NuSAP1 is required for SPB1 localization, but not its stability.

**Figure 6.**
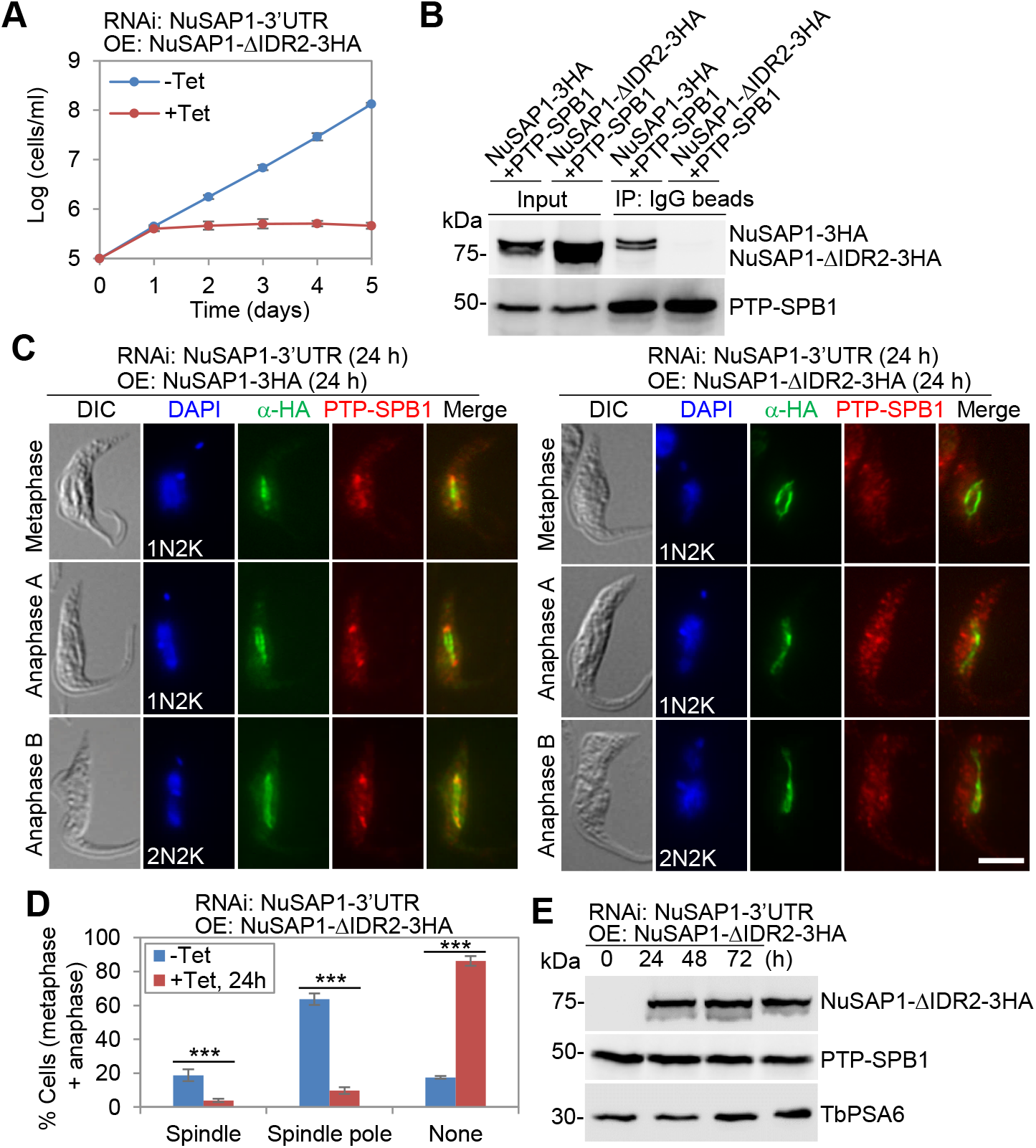
The IDR2 of NuSAP1 is required for the function of NuSAP1 and the interaction with SPB1. (**A**). The growth curve of the NuSAP1-3’UTR RNAi cell line overexpressing NuSAP1-ΔIDR2. (**B**). Co-immunoprecipitation to test the requirement of IDR2 for the interaction of NuSAP1 with SPB1. The full-length NuSAP1 served as a control. (**C**). Effect of IDR2 deletion on the localization of NuSAP1 and SPB1. Expression of the full-length NuSAP1 served as a control. Scale bar: 5 μm. (**D**). Quantitation of SPB1 localized in metaphase and anaphase cells in non-induced control and NuSAP1 RNAi/NuSAP1-ΔIDR2 OE cells. Error bars indicate S.D. from three independent experiments. ***, *p*<0.001. (**E**). Western blotting to detect the levels of endogenous PTP-tagged SPB1 and the ectopically overexpressed NuSAP1-ΔIDR2 in the NuSAP1-3’UTR RNAi/NuSAP1-ΔIDR2 OE cells. TbPSA6 served as the loading control.

### Depletion of NuSAP1 disrupts the localization of MAP103 and TbMlp2 to the spindle poles

We tested whether NuSAP1 interacts with its other proximal proteins, MAP103 and TbMlp2 (Fig. 3A). MAP103 is a spindle-associated protein with unknown function (Hayashi and Akiyoshi, 2018), and was found to be enriched at spindle poles in our previous work (Zhou and Li, 2024). TbMlp2 is a homolog of the conserved nuclear basket protein Mlp2 in yeast and humans, which is a component of the nuclear pore complex, and is enriched at spindle poles during mitosis in *T. brucei* (Holden et al., 2014; Morelle et al., 2015). Co-immunoprecipitation showed that NuSAP1 does not interact with MAP103 and TbMlp2 (Fig. 7A, B), suggesting that they localize to the proximity of NuSAP1, but do not make any direct contact with NuSAP1. Co-immunofluorescence microscopy showed that MAP103 co-localizes with NuSAP1 at the spindle and spindle poles, with MAP103 more enriched at the inner part of the spindle pole than NuSAP1 (Fig. 7C). Despite that the two proteins do not form a complex, knockdown of NuSAP1 impaired the enrichment of MAP103 at spindle poles (Fig. 7D, E), with the number of cells containing two MAP103 spindle pole foci decreased from ∼62% to ∼15% in metaphase cells and from ∼98% to ∼18% in anaphase cells and the number of cells containing no MAP103 spindle pole foci or only one MAP103 spindle pole focus increased correspondingly (Fig. 7E). These results suggest that NuSAP1 depletion likely disrupted the integrity of the spindle pole protein complex, in which one of its components NuSAP4 interacts with MAP103 and is required for MAP103 enrichment at spindle poles, thereby indirectly impairing MAP103 enrichment at spindle poles.

**Figure 7.**
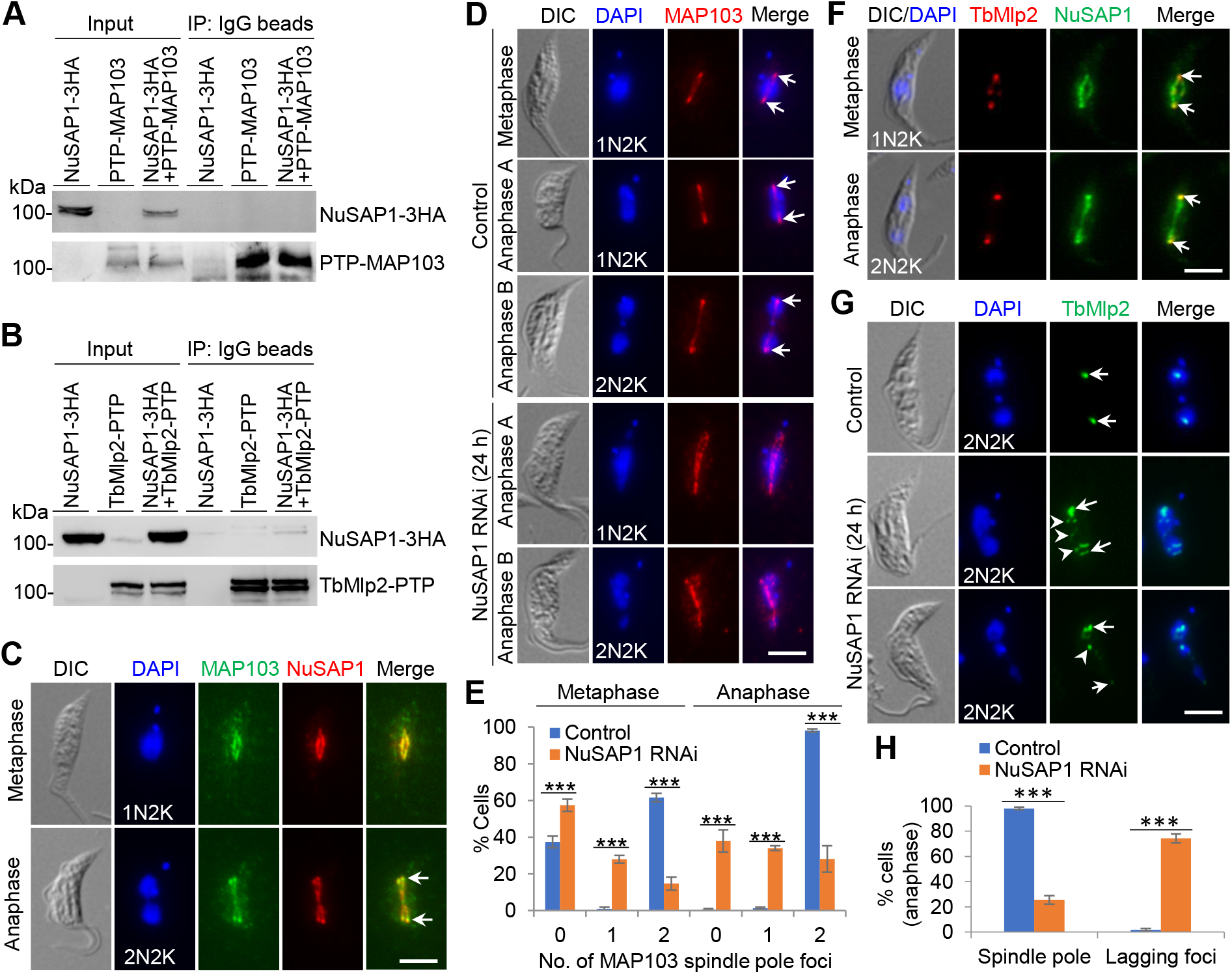
Knockdown of NuSAP1 impairs the localization of MAP103 and TbMlp2 to spindle poles. (**A**). Co-immunoprecipitation to test the *in vivo* interaction between NuSAP1 and MAP103 and between NuSAP1 and TbMlp2. (**B**). Co-localization of NuSAP1 and MAP103 to spindle and their enrichment at spindle poles. Endogenous MAP103-3HA and NuSAP1-PTP were detected by anti-HA mAb and anti-Protein A pAb, respectively. Scale bar: 5 μm. (**C**). Effect of NuSAP1 RNAi on MAP103 enrichment to spindle poles. Endogenous MAP103-PTP was detected by anti-Protein A pAb. Arrows indicate the spindle pole-enriched MAP103-PTP foci. Scale bar: 5 μm. (**D**). Quantitation of cells with different numbers of spindle pole-enriched MAP103 foci in metaphase and anaphase cells in control and NuSAP1 RNAi cells. Error bars indicate S.D. from three independent experiments. ***, *p*<0.001. (**E**). Co-localization of NuSAP1 and TbMlp2 at spindle poles. Endogenous TbMlp2-PTP and NuSAP1-3HA were detected by anti-Protein A pAb and anti-HA mAb, respectively. Arrows indicate the spindle pole-enriched foci of TbMlp2 and NuSAP1. Scale bar: 5 μm. (**F**). Effect of NuSAP1 RNAi on the localization of TbMlp2 to spindle poles. Endogenous TbMlp2-3HA was detected by anti-HA mAb. Arrows indicate TbMlp2-3HA at spindle poles, and open arrowheads indicate the lagging TbMlp2-3HA foci in NuSAP1 RNAi cells. Scale bar: 5 μm. (**G**). Quantitation of anaphase cells with lagging TbMlp2-3HA foci in control and NuSAP1 RNAi cells. Error bars indicate S.D. from three independent experiments. ***, *p*<0.001.

Immunofluorescence microscopy showed the co-localization of TbMlp2 and NuSAP1 at spindle poles (Fig. 7F), confirming TbMlp2 as a NuSAP1-proximal protein. Knockdown of NuSAP1 disrupted the localization of TbMlp2 to spindle poles, resulting in multiple lagging TbMlp2 foci (Fig. 7G) in ∼74% of the anaphase cells (Fig. 7H), in striking contrast to the exclusive spindle pole localization of TbMlp2 in ∼98% of the control cells (Fig. 7H). The enrichment of TbMlp2 at spindle poles suggests a potential interaction with certain spindle pole component(s) and a potential role in tethering the spindle pole complex into the nuclear envelope, as is in the budding yeast (Niepel et al., 2005). Thus, the disruption of the spindle pole protein complex composed of NuSAP1, NuSAP4, SPB1 and MAP103 by NuSAP1 RNAi appeared to cause the dissociation of TbMlp2 from the nuclear basket.

## Discussion

Spindle-associated proteins play important roles in regulating the establishment of a bipolar spindle for faithful chromosome segregation in eukaryotes. We previously identified a cohort of kinetoplastid-specific spindle-associated proteins and uncovered their roles in regulating chromosome segregation, but did not address the underlying mechanisms (Zhou et al., 2018; Zhou and Li, 2024). NuSAP3 interacts with the spindle microtubule depolymerase Kif13-1 and maintains Kif13-1 stability (Zhou et al., 2018), whereas NuSAP4 interacts with SPB1 and MAP103 and maintains the localization of SPB1 and MAP103 at spindle poles (Zhou and Li, 2024). NuSAP1 and NuSAP2 are both required for chromosome segregation (Zhou et al., 2018), but their mechanistic roles remained elusive. We reported here that NuSAP1 is capable of bundling microtubules *in vitro* (Fig. 1) and is required for bipolar spindle assembly *in vivo* (Fig. 2). NuSAP4 is also required for bipolar spindle assembly *in vivo* (Zhou and Li, 2024), but it does not bundle microtubules *in vitro* (Fig. 1). NuSAP1 is thus the first spindle-associated protein in *T. brucei* found to be capable of bundling microtubules. Structurally, NuSAP1 does not appear to contain any conserved microtubule-binding domains, but two blocks of basic residues (a.a. 354-371 and a.a. 689-703) might contribute to microtubule binding, as is the block of basic residues in the kinetochore protein KKT4 (Llauro et al., 2018), although this needs to be experimentally verified. The mechanism underlying the microtubule-bundling activity of NuSAP1 remains unclear, but we hypothesize that NuSAP1 may form a multimer, so the oligomerization of NuSAP1 brings multiple microtubules together to form microtubule bundles. Alternatively, NuSAP1 may contain multiple microtubule-binding domains, which bind different microtubules, thereby bringing multiple microtubules together to form microtubule bundles. No matter which mechanism NuSAP1 employs to bundle microtubules, the *in vitro* activity and the spindle localization suggest its cellular function in bundling spindle microtubules to promote bipolar spindle assembly.

The defective assembly of a bipolar spindle in NuSAP1 RNAi cells (Fig. 2) is in line with the *in vitro* microtubule-bundling activity of NuSAP1 and the localization of NuSAP1 to the spindle (Fig. 1). How the microtubule-bundling activity of NuSAP1 contributes to bipolar spindle assembly correlates with the structural organization of the spindle. In a typical mitotic spindle, three types of microtubules are present; kinetochore microtubules that capture kinetochores, interpolar microtubules that maintain spindle bipolarity and overall structure, and astral microtubules that position the spindle and maintain spindle stability. While astral microtubules exist as individual filament, kinetochore microtubules and interpolar microtubules often organize as bundles (Pavin and Tolic, 2016). Kinetochore microtubule bundles (or kinetochore fibers) consist of parallel microtubules emanated from the same spindle pole, whereas interpolar microtubule bundles consist of anti-parallel microtubules emanated from opposite spindle poles, which overlap at the central portion of the spindle(Cheeseman and Desai, 2008; Mastronarde et al., 1993). In *T. brucei*, astral microtubules likely do not exist, but bundles of kinetochore microtubules and interpolar microtubules are readily detectable (Ogbadoyi et al., 2000). NuSAP1 may contribute to the bundling of both kinetochore microtubules and interpolar microtubules to promote the establishment of a bipolar spindle. In the absence of NuSAP1 due to RNAi-mediated knockdown, spindle microtubules failed to form bundles, thereby disrupting the typical spindle shape (Fig. 2).

Although NuSAP1 is capable of bundling microtubules *in vitro* without any co-factors, it forms a complex *in vivo* in trypanosome cells with NuSAP4, which bears structural resemblance to, and binds to, the CC2 motif of NuSAP1 (Figs. 3 and 4). The interdependency between NuSAP1 and NuSAP4 for their respective protein stability in trypanosome cells suggests that the NuSAP1-NuSAP4 complex may serve as a single functional unit, in which the loss of either protein destabilizes the unit. Knockdown of NuSAP4 exerts weaker effects on NuSAP1 stability than the knockdown of NuSAP1 on NuSAP4 stability (Fig. 3H), which could be attributed to the lesser efficiency of NuSAP4 knockdown (Fig. 3H). Due to its small size and lack of microtubule-bundling activity, NuSAP4 might simply function to stabilize NuSAP1 in the unit for the latter to fulfil the microtubule-bundling activity to promote bipolar spindle assembly. However, loss of the NuSAP4-binding CC2 motif disrupted NuSAP1 function despite its localization to the spindle (Fig. 4), implying that NuSAP4 and its binding to NuSAP1 have additional functions, which remain to be explored. Nonetheless, the destabilization of NuSAP4 in cells expressing NuSAP1-ΔCC2 further confirmed that the interaction with NuSAP1 stabilizes NuSAP4.

NuSAP1 also forms a complex with the spindle-localized and spindle pole-enriched protein SPB1, but not the spindle-localized and spindle pole-enriched protein MAP103 (Figs. 5 and 7). NuSAP4, however, interacts with both SPB1 and MAP103, albeit having weaker association with the latter (Zhou and Li, 2024). Since NuSAP4 can pull down NuSAP1 (Fig. 3) and MAP103 (Zhou and Li, 2024), the inability for NuSAP1 to pull down MAP103 suggests that the three proteins likely form separate protein complexes, the NuSAP1-NuSAP4 complex and the NuSAP4-MAP103 complex, or a tri-protein complex with MAP103 transiently or loosely associating with NuSAP4. This notion is reflected by the least detected peptides of MAP103 among the NuSAP1-proximal proteins identified by BioID (Fig. 3A). In contrast, both NuSAP1 (Fig. 5) and NuSAP4 (Zhou and Li, 2024) are able to pull down SPB1, suggesting that these three proteins may form a tri-protein complex, with NuSAP4 and SPB1 interacting with distinct domains of NuSAP1 (Figs. 4 and 6), although we cannot rule out the possibility that they form three separate protein complexes, the NuSAP1-NuSAP4 complex, the NuSAP1-SPB1 complex, and the NuSAP4-SPB1 complex. However, the dependency of NuSAP4 on NuSPA1 for stability (Fig. 3F) argues against the presence of the NuSAP4-SPB1 complex, in which this portion of NuSAP4 should not be affected by NuSAP1 RNAi. Same is true for the NuSAP4-MAP103 complex, which is unlikely to exist. Given that SPB1 and MAP103 do not interact *in vivo* (Zhou and Li, 2024), it is more likely that these four proteins may form two separate tri-protein complexes, the NuSAP1-NuSAP4-SPB1 complex and the NuSAP1-NuSAP4-MAP103 complex, despite that in the latter complex MAP103 only loosely or transiently associates with NuSAP4.

The enrichment of NuSAP1 at spindle poles (Fig. 1) raises a question of whether NuSAP1, with a microtubule-bundling activity, catalyzes microtubule bundling at spindle poles. Despite the lack of a canonical centriole structure at the spindle pole in trypanosomes, a ring-shaped structure located at the spindle pole is detectable, and multiple microtubules converge into a single focus that is connected to the nuclear envelope (Ogbadoyi et al., 2000), suggestive of a putative microtubule-organizing center at the spindle pole. It is unclear whether the convergence of microtubules at the spindle pole is attributed to NuSAP1’s microtubule bundling activity, or the emanation of multiple microtubules from the single spindle pole shows an appearance of microtubule convergence. In the latter scenario, NuSAP1 may play an additional role at the spindle pole by assembling the spindle pole protein complex composed of NuSAP1, NuSAP4, SPB1, and MAP103.

The disruption of TbMlp2 location at the spindle pole by NuSAP1 depletion (Fig. 7) raises a question of how the nuclear basket of the nuclear pole complex may associate with the spindle pole in trypanosomes. The enrichment of TbMlp2, a conserved nuclear basket protein of the nuclear pore complex, at spindle poles and the role of TbMlp2 in regulating chromosome segregation (Holden et al., 2014; Morelle et al., 2015) suggest that the spindle pole is likely anchored to the nuclear basket of the nuclear pore complex or that the nuclear basket determines the position of the spindle pole. Although co-immunoprecipitation does not detect *in vivo* interactions between TbMlp2 and NuSAP1 (Fig. 7) and between TbMlp2 and NuSAP4 (Zhou and Li, 2024), BioID with NuSAP1 as bait appeared to have detected large amounts of TbMlp2 protein, as reflected by the large numbers of TbMlp2 peptides detected by mass spectrometry (Fig. 3A), suggesting the very close proximity of TbMlp2 to NuSAP1. The dependency of TbMlp2 localization on NuSAP1 (Fig. 7) and NuSAP4 (Zhou and Li, 2024) suggests the requirement of an intact NuSAP1- and NuSAP4-containing spindle pole protein complex for TbMlp2 localization. However, given the lack of a direct interaction between TbMlp2 and the NuSAP1-NuSAP4 complex, it is possible that the disruption of the spindle pole complex by NuSAP1 RNAi and NuSAP4 RNAi destabilizes the association of the nuclear basket with the spindle pole, leading to the dissociation of TbMlp2 from the nuclear basket. Alternatively, TbMlp2 may interact with an unidentified spindle pole component that depends on the NuSAP1-NuSAP4 complex, whereby mediating the association between the nuclear basket and the spindle pole, as is the case of the budding yeast Mlp2p protein that interacts with three spindle pole components (Niepel et al., 2005). The structural organization and the molecular composition of the spindle pole in *T. brucei* remain unresolved, but they are believed to be different from that in yeast and animals due to the absence of a canonical centriole and the conserved core components of the centrosome at the spindle pole. Moreover, how the spindle pole may physically connect to the nuclear basket of the nuclear pore complex in *T. brucei* is also elusive. Thus, further exploration of the spindle pole, the nuclear basket, and how they may physically associate may provide new insights into the mechanisms of bipolar spindle assembly in this early branching unicellular eukaryote.

## Materials and Methods

### Prediction of protein and protein complex structures by AlphaFold3

The structures of NuSAP1, the NuSAP1-NuSAP4 complex, the CC2^NuSAP1^-NuSAP4 complex, the NuSAP1-NuSAP4-SPB1 complex were predicted by AlphaFold3 (Abramson et al., 2024) using the webserver https://alphafoldserver.com/. The predicted structures were visualized and exported with the online Mol* 3D viewer at the following webserver: https://www.rcsb.org/3d-view.

### Trypanosome cell culture and RNA interference

The 29-13 strain of the procyclic form of *T. brucei* (Wirtz et al., 1999) was cultured in SDM-79 medium supplemented with 10% heat-inactivated fetal bovine serum (MilliporeSigma), 15 µg/ml G418, and 50 µg/ml hygromycin, and the Lister427 strain of the procyclic form of *T. brucei* was cultured in SDM-79 medium supplemented with 10% fetal bovine serum at 27°C. The NuSAP1 RNAi cell line (Zhou et al., 2018) and the NuSAP4 RNAi cell line (Zhou and Li, 2024) were cultured in SDM-79 medium supplemented with 10% heat-inactivated fetal bovine serum, 15 µg/ml G418, 50 µg/ml hygromycin, and 2.5 µg/ml phleomycin at 27°C. To induce RNAi, the RNAi cell lines were incubated with 1.0 µg/ml tetracycline.

### *In situ* epitope tagging of proteins

Tagging of proteins from their endogenous locus with an epitope at either the N-terminus or the C-terminus was performed using the PCR-based method reported previously (Shen et al., 2001).

For co-immunofluorescence microscopy of the co-localization of NuSAP1 with β-tubulin, Kif13-1, TbMlp2, SPB1, and MAP103, NuSAP1 was endogenously tagged with a C-terminal PTP epitope and the other proteins were endogenously tagged with a C-terminal triple HA epitope in the Lister427 strain. For co-immunofluorescence microscopy of the co-localization of NuSAP1 and NuSAP4, NuSAP1 was endogenously tagged with a C-terminal triple HA epitope and NuSAP4 was endogenously tagged with a C-terminal PTP epitope in the Lister427 strain.

For co-immunoprecipitation analysis of the interaction between NuSAP1 and NuSAP4, the interaction between NuSAP1 and MAP103, and the interaction between NuSAP1 and TbMlp2, NuSAP1 was endogenously tagged with a C-terminal triple HA epitope, NuSAP4 and TbMlp2 were endogenously tagged with a C-terminal PTP epitope, and MAP103 was endogenously tagged with an N-terminal PTP epitope in the Lister427 strain. For co-immunoprecipitation analysis of the interaction between NuSAP1 and SPB1, NuSAP1 was endogenously tagged with a C-terminal PTP epitope, and SPB1 was endogenously tagged with a C-terminal triple HA epitope in the Lister427 strain.

For immunofluorescence microscopy of spindle assembly in NuSAP4 RNAi cells, KIN-F was endogenously tagged with a C-terminal triple HA epitope, and MAP103 was endogenously tagged with an N-terminal PTP epitope in the NuSAP1 RNAi cell line. For immunofluorescence microscopy of the effect of NuSAP1 RNAi on the localization of NuSAP4, SPB1, MAP103, and TbMlp2, NuSAP4, SPB1, and TbMlp2 were each tagged with a C-terminal triple HA epitope and MAP103 was tagged with an N-terminal PTP epitope in NuSAP1 RNAi cell line. For immunofluorescence microscopy of the effect of NuSAP4 RNAi on NuSAP1, NuSAP1 was endogenously tagged with a C-terminal triple HA epitope in the NuSAP4 RNAi cell line.

Transfectants were selected with appropriate antibiotics, and clonal cell lines were obtained by limiting dilution in a 96-well plate containing SDM-79 medium supplemented with 20% fetal bovine serum and appropriate antibiotics.

### Generation of NuSAP1 RNAi complementation cell lines

To generate NuSAP1 RNAi complementation cell lines, a NuSAP1-3’UTR RNAi cell line was first generated by cloning a 575-bp fragment of the 3’UTR of NuSAP1 into the pZJM vector. The resulting plasmid, pZJM-NuSAP1-3’UTR, was used to electroporate the 29-13 strain, and transfectants were selected with 2.5 µg/ml phleomycin. Clonal cell lines were generated by limiting dilution as described above. To monitor RNAi efficiency, NuSAP1 was endogenously tagged with an N-terminal PTP epitope in the NuSAP1-3’UTR RNAi cell line by PCR-based method reported previously (Shen et al., 2001) and described above.

The full-length *NuSAP1* gene and the mutant *NuSAP1* gene lacking the sequence encoding the CC2 or the IDR2 were cloned into the pLew100-3HA-PAC vector. The resulting plasmids, pLew100-NuSAP1-3HA-PAC, pLew100-NuSAP1-ΔCC2-3HA-PAC, and pLew100-NuSAP1-ΔIDR2-3HA-PAC, were used to electroporate the NuSAP1-3’UTR RNAi cell line. Successfully transfectants were selected with 1.0 µg/ml puromycin and cloned by limiting dilution as described above. To induce NuSAP1-3’UTR RNAi and ectopic expression of NuSAP1-3HA, NuSAP1-ΔCC2-3HA, and NuSAP1-ΔIDR2-3HA, cells were incubated with 1.0 μg/ml tetracycline, and cell growth was monitored daily.

Using the PCR-based protein epitope-tagging method (Shen et al., 2001), NuSAP4 was endogenously tagged with a C-terminal PTP epitope in the NuSAP1-3’UTR RNAi/NuSAP1-3HA OE cell line and the NuSAP1-3’UTR RNAi/NuSAP1-ΔCC2-3HA OE cell line, and SPB1 was endogenously tagged with an N-terminal PTP epitope in the NuSAP1-3’UTR RNAi/NuSAP1-ΔIDR2-3HA OE cell line and the NuSAP1-3’UTR RNAi/NuSAP1-3HA OE cell line. Transfectants were selected with 10 μg/ml blasticidin and cloned by limiting dilution. Cells were induced with 1.0 μg/ml tetracycline and used for immunofluorescence microscopy, western blotting, and co-immunoprecipitation. Since NuSAP4-PTP level was reduced in NuSAP1-3’UTR RNAi cells, 10 μg/ml MG-132 was added to cells for 8 hours to prevent NuSAP4-PTP degradation before performing co-immunoprecipitation.

### Co-immunoprecipitation and western blotting

Cells co-expressing endogenously 3HA-tagged NuSAP1 and PTP-tagged NuSAP4, MAP103, or TbMlp2, cells co-expressing endogenously PTP-tagged NuSAP1 and 3HA-tagged SPB1, and cells co-expressing endogenously PTP-tagged NuSAP4 and ectopically overexpressed, 3HA-tagged NuSAP1 or NuSAP1-ΔCC2 were lysed in 1.0 ml immunoprecipitation buffer (25 mM Tris-HCl, pH7.6, 100 mM NaCl, 1 mM DTT, 1% NP-40, and protease inhibitor cocktail) for 30 min on ice. Cell lysate was centrifugated at the highest speed in a microcentrifuge at 4°C, and cleared cell lysate was incubated with 50 µl IgG beads for 1 h at 4°C with gentle rotation. The immunoprecipitate was washed six times with the immunoprecipitation buffer, and bound proteins were eluted by boiling with the 1x SDS sampling buffer. Eluted proteins were separated by SDS-PAGE, transferred onto a PVDF membrane, and immunoblotted with the anti-HA antibody and the anti-Protein A antibody. Cells expressing NuSAP1-3HA alone, NuSAP1-PTP alone, NuSAP4-PTP alone, SPB1-3HA alone, PTP-MAP103 alone, and TbMlp2-PTP alone were included as negative controls.

### Immunofluorescence microscopy

Trypanosome cells were briefly washed with PBS, adhered to the glass coverslips, fixed with cold methanol (-20°C), and then rehydrated with PBS. Cells on the coverslips were blocked with 3% BSA in PBS, and incubated with the KMX-1 antibody (anti-β-tubulin monoclonal antibody), FITC-conjugated anti-HA monoclonal antibody for 3HA-tagged proteins (1:400 dilution, Millipore-Sigma) and/or anti-Protein A polyclonal antibody for PTP-tagged proteins (1:400 dilution, Millipore-Sigma). The coverslips were washed three times with PBS, and incubated with Cy3-conjugated anti-mouse IgG (1:400 dilution, Millipore-Sigma) to detect β-tubulin or Cy3-conjugated anti-rabbit IgG (1:400 dilution, Millipore-Sigma) to detect PTP-tagged proteins. The coverslips were washed three times with PBS and mounted with DAPI-containing VectaShield mounting medium (Vector Labs). Cells on the coverslips were imaged under an inverted fluorescence microscope (Olympus IX71) equipped with a cooled CCD camera (model Orca-ER, Hamamatsu). Images were acquired using the Slidebook software.

### Expression and purification of recombinant proteins

The full-length coding sequences of the *NuSAP1* gene, the *NuSAP4* gene, and the *TbSpef1* gene were each amplified from the genomic DNA by PCR and cloned into the pGEX-4T-3 vector to express GST-fused NuSAP1, NuSAP4, and TbSpef1. The resultant plasmids were transformed into the *E. coli* strain BL-21, and the bacterial cells were induced to express the recombinant protein with IPTG for 16 h at 16°C. Cells expressing the recombinant protein were lysed by sonication, and the cell lysate was cleared by centrifugation at 20,627 *g* for 10 min at 4°C. The cleared cell lysate was then incubated with the Glutathione Sepharose 4B beads for 1 h at 4°C with gentle agitation. The Glutathione Sepharose 4B Beads were washed three times with 0.1% Triton X-100 in PBS, and bound proteins were eluted with 20 mM glutathione in 50 mM Tris, pH 9.0, 0.1% Triton X-100, 100 mM NaCl, and 1 mM DTT. Eluted proteins were concentrated, and the buffer was exchanged with Amicon Ultra Centrifugal Filters 10K (Millipore-Sigma).

### Tubulin polymerization and microtubule-bundling assay

*In vitro* tubulin polymerization using porcine brain tubulin was performed using the protocols described previously (Hu et al., 2024). Non-labeled porcine brain tubulin (Cytoskeleton, Inc., Cat#: T240-A80, 80 μg) was mixed with rhodamine-labeled porcine brain tubulin (Cytoskeleton, Inc., Cat#: TL590M, 20 μg) in the BRB80-DTT buffer (80 mM Potassium-PIPES, pH 6.8, 1.0 mM MgCl_2_, 1.0 mM EGTA, 1.0 mM DTT) supplemented with 1.0 mM Guanylyl-(α,β)-methylene-diphosphonate (GMP-CPP), and the mixture was incubated at 4°C for 5 min. The mixture was centrifugated at 279,000 ×*g* for 5 min at 4°C in a TLA120.1 rotor in an ultracentrifuge (Beckman Coulter TL-100), and the supernatant was snap-frozen as aliquots in liquid nitrogen and stored at -80°C. To make microtubule seeds, an aliquot of the supernatant was 1:4 diluted with the BRB80-DTT buffer and incubated at 37°C for 30 min. Microtubule seeds were pelleted at 353,000 ×*g* at 27°C for 5 min and resuspended in the BRB80-DTT buffer. To polymerize microtubules from the microtubule seeds, rhodamine-labeled tubulin was mixed with the microtubule seeds in the BRB80-DTT buffer supplemented with 2.0 mM GTP. Microtubules were then successively assembled by incubating with 1 μl of 1 μM Taxol in BRB80-DTT-GTP buffer (BRB80-DTT buffer plus 1.0 mM GTP) for 20 min, 1.1 μl of 10 μM Taxol for 10 min, and 1.2 μl of 100 μM Taxol for 10 min at 37°C. For the microtubule-bundling assay, the assembled microtubules were centrifuged at 1,500 ×*g* for 3 min to remove any Taxol-induced microtubule bundles. Purified recombinant proteins were centrifuged at 279,000 ×*g* for 15 min at 4°C in a TLA120.1 rotor in an ultracentrifuge (Beckman Coulter TL-100) to remove protein aggregates. Microtubules were incubated with 100 nM purified recombinant proteins in the BRB80-DTT-GTP buffer containing 10 μM Taxol, 5% glycerol, and 1.0 mM ATP at room temperature for up to 3 minutes. Time-course samples (20 sec, 60 sec, and 3 min) were collected and adhered onto glass coverslips. Microtubules were fixed with 4% paraformaldehyde and imaged under a fluorescence microscope.

## Author contributions

**Qing Zhou:** Conceptualization, Methodology, Visualization, Validation, Investigation, Formal analysis, Writing – Reviewing and Editing. **Yasuhiro Kurasawa**: Methodology, Visualization, Validation, Investigation, Formal analysis. **Thiago Souza Onofre:** Methodology, Visualization, Validation, Investigation, Formal analysis. **Ziyin Li**: Conceptualization, Supervision, Project administration, Funding acquisition, Writing – Original Draft, Writing – Reviewing and Editing.

## Acknowledgements

This work was supported by the NIH R01 grants AI118736 and AI101437 to Z. L.

## Competing interests

The authors declare that they have no conflicts of interest with the contents of this article.

## Data availability

All data are contained within the manuscript.

## Notes

### Competing Interest Statement

The authors have declared no competing interest.

